# Integrating cross-linking experiments with *ab initio* protein-protein docking

**DOI:** 10.1101/275891

**Authors:** Thom Vreven, Devin K. Schweppe, Juan D. Chavez, Chad R. Weisbrod, Sayaka Shibata, Chunxiang Zheng, James E. Bruce, Zhiping Weng

## Abstract

*Ab initio* protein-protein docking algorithms often rely on experimental data to identify the most likely complex structure. We integrated protein-protein docking with the experimental data of chemical cross-linking followed by mass spectrometry. We tested our approach using 12 cases that resulted from an exhaustive search of the Protein Data Bank for protein complexes with cross-links identified in our experiments. We implemented cross-links as constraints based on Euclidean distance or void-volume distance. For most test cases the rank of the top-scoring near-native prediction was improved by at least two fold compared with docking without the cross-link information, and the success rates for the top 5 and top 10 predictions doubled. Our results demonstrate the delicate balance between retaining correct predictions and eliminating false positives. Several test cases had multiple components with distinct interfaces, and we present an approach for assigning cross-links to the interfaces. Employing the symmetry information for these cases further improved the performance of complex structure prediction.

**Highlights:**

- Incorporating low-resolution cross-linking experimental data in protein-protein docking algorithms improves performance more than two fold.
- Integration of protein-protein docking with chemical cross-linking reveals information on the configuration of higher order complexes.
- Symmetry analysis of protein-protein docking results improves the predictions of multimeric complex structures

## INTRODUCTION

Protein interactions play critical roles in biological processes, including the immune system, signaling pathways, and enzymatic reactions. Proteome-wide studies have shown that most proteins interact with one or more other proteins [1]. Three-dimensional structures of protein-protein complexes are needed to understand these processes, which can be carried out at the atomic resolution by X-ray crystallography, nuclear magnetic resonance, or cryoelectron microscopy. But these experiments are difficult to perform and sometimes do not succeed in determining the structures.

Various other experimental techniques can provide structural information at lower resolution. H/D exchange, mutagenesis experiments (in particular alanine scanning), and chemical cross-linking followed by mass spectrometry can identify interfacial residues or residue pairs, while small-angle X-ray scattering (SAXS) and electron microscopy can provide orientational information that is not residue specific [2]. A number of computational methods have been developed to predict protein-protein complex structures, but typically yielding many incorrect predictions—if a computational algorithm is allowed to make ten predictions for a protein-protein complex, it has roughly a fifty percent chance to yield at least one near-native structure [3-6]. Integrating computational algorithms with lower-resolution experimental data can improve the accuracy of protein complex structures prediction [7-15]. The experimental data can either be used to guide computational prediction [16,17] or to filter predictions in a post-processing step [11].

In this study, we integrated the *ab initio* protein-protein docking algorithm ZDOCK [18-21] with the experimental data of chemical cross-linking followed by mass spectrometry. Cross-linking reagents can form covalent bonds with protein residues that are closer in distance than the length of the linker. Trypsin digestion of cross-linked proteins, followed by mass spectrometry, identifies protein residues that were cross-linked. The cross-linking reagent has a maximum length; therefore, the cross-linking data give an upper bound for the geometric distance between paired residues. Cross-linking data has been used extensively to validate or guide protein-protein docking predictions [11,22-25], and various approaches were developed to integrate the constraints with the docking algorithms [11,26-28]. Systematic investigations of the performance using large data sets were, however, carried out only using simulated cross-linking data [27]. Here we present a data set that is derived from our proteome-wide experiments [29-34] and all use the same linker. The dataset was searched against the known structures in the Protein Data Bank [35] and yielded 12 test cases. Although the resulting collection of test cases is limited in size, it enabled us to compare the effectiveness of several integration schemes and develop a new algorithm for associating the cross-links with specific interfaces in higher-order protein-protein complexes.

## RESULTS AND DISCUSSION

### Overall approach

We used ZDOCK [18-21] with input component proteins obtained from X-ray crystallography or through homology modeling using X-ray crystallography template structures. The ZDOCK algorithm was integrated with experimental cross-linking data to generate only predictions that satisfy the cross-links. The following three approaches were tested: (1) Filtering the predictions from a standard ZDOCK calculation using the Euclidean distance between cross-linked sites. Although Euclidean distances are fast to compute and therefore applicable to large sets of predictions, the cross-linking distances could be underestimated because the Euclidean path is allowed to pass through protein-occupied space. (2) Filtering the ZDOCK predictions using the Xwalk algorithm [27,36] to determine the shortest path that is allowed to only pass through protein-unoccupied space (void-volume). Although physically more accurate than Euclidean distances, computationally the grid-based algorithm is orders of magnitude more expensive to evaluate. (3) Restrict ZDOCK to search only the space that satisfies the Euclidean cross-linking constraints. This approach yields more retained predictions than the filtering methods and therefore may improve performance.

We performed cross-linking and mass spectrometry experiments with the lysine-reactive BDP-NHP chemical [30] and then used the ReACT [30] algorithm to identify the cross-linked sites. We used our previously published cross-linking data [29-34] and unpublished data. We only analyzed heteromeric interactions in this study because most *ab initio* protein-docking algorithms are designed to predict such complexes, and we plan to investigate homomeric complexes in future studies. Using the cross-linking data, we searched the Protein Data Bank (PDB) [35] and retained 12 complexes that had matching unbound component proteins and are thus suitable for testing rigid-body docking algorithms such as ZDOCK. In this search we used a sequence identity cutoff of 30%, and used homology modeling to generate the structures if needed. We used the resulting collection of complexes to assess the three approaches for integrating cross-linking experiments and protein-protein docking algorithms.

### Test set

Our test set contains 12 protein-protein complexes. Ideally, a protein-protein docking test set has the unbound structures available for all component proteins, or unbound templates that can be used for homology modeling the components. However, in order to maximize the number of entries in our set, we allowed seven tests that had one of the components in the bound form. One of these cases had the unbound structure for the other component available (*unbound/bound docking*), and for the remaining six the other component needed to be homology modeled (*homology/bound docking*). For one case in our set we had unbound structures for both components (*unbound/unbound docking*), for three cases only unbound templates (*homology/homology docking*), and for the final case one template and one unbound structure (*homology/unbound docking*). The 12 complexes could be divided into eight groups based on fold-similarity and are summarized in Table 1 and shown in Figure 1.

**Table 1:** The test set.

| Case<sup>1</sup> | UniProt 1 | UniProt 2 | Bound PDB<sup>2</sup> | Unbound PDB 1<sup>3</sup> | Unbound PDB 2<sup>4</sup> | I-RMSD<sup>5</sup> |
| --- | --- | --- | --- | --- | --- | --- |
| **1**<br>Tubulin | β-subunit<br>P04350 | α-subunit<br>P68363 | 4X20<br>(97%/100%) | 1Z5V<br>(34%) | 1Z5V<br>(32%) | 1.95 (1.88/2.02) [B:A]<br>2.24 (2.25/2.24) [B:C] |
| **2A**<br>Succinyl-CoA ligase | α-subunit<br>P53597 | β-subunit<br>Q96I99 | 1EUC<br>(96%/96%) | 1OI7<br>(54%) | none | 1.02 (1.53/ n/a) |
| **2B**<br>Succinyl-CoA ligase | α-subunit<br>P0AGE9 | β-subunit<br>P0A836 | 1SCU<br>(100%/100%) | 1OI7<br>(44%) | none | 0.66 (1.02/ n/a) |
| **2C**<br>Succinyl-CoA ligase | α-subunit<br>B7I6T1 | β-subunit<br>B7I6T2 | 1SCU<br>(71%/65%) | 1OI7<br>(57%) | none | 3.56 (5.19/ n/a) |
| **2D**<br>Succinyl-CoA ligase | α-subunit<br>Q51567 | β-subunit<br>P53593 | 1SCU<br>(89%/77%) | 1OI7<br>(58%) | none | 2.72 (3.95/ n/a) |
| **3A**<br>F1-ATP synthase | β-subunit<br>P06576 | α-subunit<br>P25705 | 1COW<br>(99%/98%) | 4Q4L<br>(70%) | 2R9V<br>(59%) | 4.72 (5.13/4.29) [A:D]<br>5.06 (6.05/4.14) [A:E] |
| **3B**<br>F1-ATP synthase | β-subunit<br>P0ABB4 | P0ABB0 | 3OAA<br>(100%/100%) | 4Q4L<br>(80%) | 2R9V<br>(55%) | 4.76 (5.30/4.21) [A:D]<br>4.79 (5.65/3.87) [A:E] |
| **4**<br>Profilin- β-actin complex | Profilin-1<br>P07737 | β-actin<br>P60709 | 2BTF<br>(95%/100%) | 1PNE<br>(95%) | 1IJJ (94%) | 0.75 (0.40/0.99) |
| **5**<br>Alu binding proteins | SRP14<br>P37108 | SRP9<br>P49458 | 4UYK<br>(100%/100%) | 2W9J<br>(31%) | none | 1.51 (2.12/ n/a) |
| **6**<br>BAM complex | BamD<br>P0AC02 | BamC<br>P0A903 | 3TGO<br>(100%/100%) | 3Q5M<br>(100%) | none | 1.36 (1.78/ n/a) |
| **7**<br>GTPase | Inhibitor<br>P50395 | Rab14<br>P61106 | 1UKV<br>(56%/46%) | 1GND<br>(87%) | 4D0G<br>(99%) | 2.20 (1.56/3.02) |
| **8**<br>ElonginB- ElonginC | ElonginC<br>Q15369 | ElonginB<br>Q15370 | 1VCB<br>(100%/100%) | 1HV2<br>(40%) | none | 1.46 (1.99/ n/a) |

<sup>1</sup> Human proteins are listed in roman, and other proteins in italic.
<sup>2</sup> In parentheses are the sequence identities between the PDB entries and the UniProt sequences in the second and third columns.
<sup>3</sup> In parentheses are the sequence identities between the PDB entries and the UniProt 1 sequences in the second column.
<sup>4</sup> In parentheses are the sequence identities between the PDB entries and the UniProt 2 sequences in the third column.
<sup>5</sup> In parentheses are the I-RMSDs for the two individual proteins. When an unbound structure was not available, the I-RMSD became formally zero and is listed as n/a. I-RMSDs are given for distinct interfaces when present, which are denoted in square brackets.

**Figure 1:**
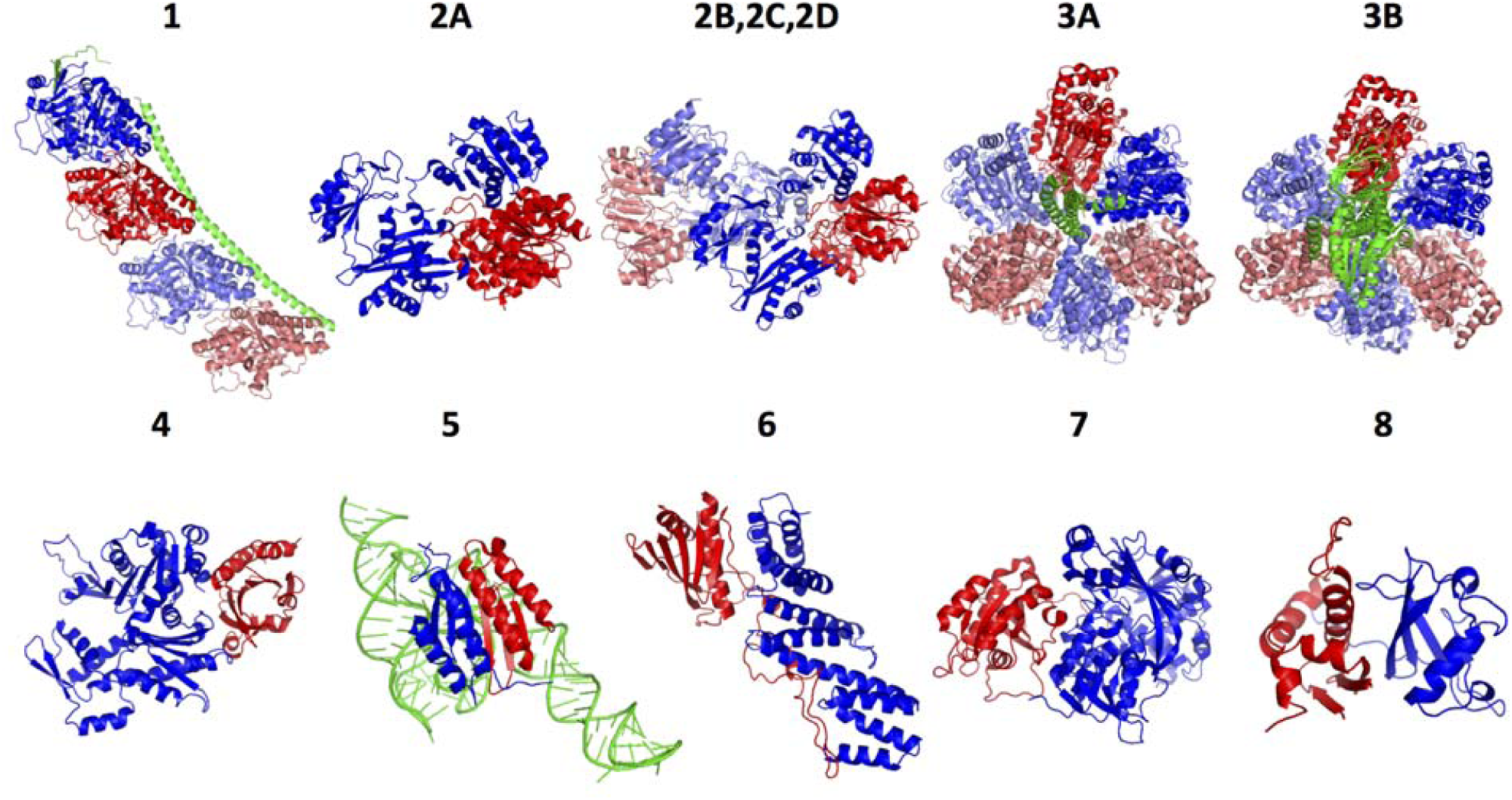
Complexes in the test set. Unique components are in red and blue, and the equivalent (same sequence) components in light red and light blue. The green protein chains in **1**, **3A**, and **3B**, and RNA in **5** were not included in the prediction and analysis.

The I-RMSD values in Table 1 indicate the conformational differences between the bound and unbound interfaces [6]. Except for **2C**, **2D**, **3A**, and **3B** that had I-RMSDs ranging from 3 to 5 Å, the other complexes had I-RMSDs under 2.5 Å and would be classified as having low to medium difficulty for *ab initio* docking algorithms such as ZDOCK [6]. Table 2 lists the experimentally detected cross-links. The cross-linked proteins had 1-7 pairs of cross-linked sites, averaging just under 3 per protein pair. Based on the bound structures, the Euclidean distances between the cross-linked sites were under 35 Å and the void-volume distances under 40 Å, except for cases **2C**, **2D**, and **6**, whose cross-linked sites were at slightly greater distances. The distance distribution is consistent with previously reported data [29].

**Table 2:** Cross-linked sites for the test cases with the Euclidean and void-volume distances (using C_β_ atoms) computed using the bound structures (for each case ordered by Euclidean distance).

| Case | Residue 1 | Residue 2 | Euclidean distance (Å) | Void-volume distance (Å) |
| --- | --- | --- | --- | --- |
| **1** | 216 | 326 | 14.4 | 16.9 |
|  | 216 | 336 | 22.1 | 25.0 |
|  | 336 | 394 | 22.3 | 30.6 |
|  | 324 | 394 | 24.7 | 28.5 |
|  | 216 | 338 | 27.7 | 31.3 |
| **2A** | 192 | 403 | 26.0 | 29.9 |
|  | 90 | 271 | 28.0 | 32.0 |
| **2B** | 256 | 146 | 15.3 | 15.5 |
|  | 241 | 56 | 18.2 | 20.1 |
|  | 285 | 372 | 20.5 | 23.1 |
|  | 144 | 372 | 22.2 | 24.9 |
|  | 256 | 56 | 22.8 | 27.3 |
|  | 144 | 359 | 26.6 | 28.6 |
|  | 43 | 172 | 33.8 | 37.5 |
| **2C** | 43 | 173 | 35.4 | 41.0 |
| **2D** | 43 | 175 | 39.0 | 44.2 |
| **3A** | 133 | 123 | 14.7 | 18.9 |
|  | 480 | 427 | 20.1 | 25.3 |
|  | 522 | 427 | 20.9 | 23.3 |
|  | 522 | 531 | 24.6 | 25.3 |
|  | 485 | 506 | 25.3 | 28.5 |
|  | 480 | 531 | 26.4 | 31.4 |
|  | 519 | 531 | 26.6 | 28.3 |
| **3B** | 372 | 419 | 16.4 | 21.2 |
| **4** | 70 | 291 | 15.5 | 17.9 |
|  | 54 | 113 | 23.7 | 27.0 |
| **5** | 66 | 30 | 14.8 | 20.2 |
|  | 19 | 30 | 15.6 | 26.7 |
| **6** | 149 | 166 | 38.8 | 42.5 |
| **7** | 221 | 61 | 31.1 | 34.0 |
| **8** | 72 | 46 | 22.3 | 27.2 |

Three of the groups (1-3) display multiple interfaces in the bound structures. When they involved different binding sites they were considered both separately and combined in the assessment of the prediction algorithms. The first example, group 1, consists of the complex of tubulin α and β chains (**1**). The bound structure contains a stathmin chain as well, although it was not incorporated in the docking experiments [37]. The structure shows three interfaces between the components, two of which are distinct and involve two different binding sites for each component.

Group 2 consists of four members, human (**2A**) and non-human (**2B**, **2C**, **2D**) Succinyl-CoA ligases, each being a complex of a α-subunit and a β-subunit. The β-subunit consists of two domains that both bind the α-subunit. The sequence identities amongst the four members of the group range from 44% to 89% (the latter being between the α-subunits of **2B** and **2D**), and we considered these complexes as distinct test cases for two reasons. First, there is little redundancy among the cross-links (only the single cross-link of **2C** was equivalent to one of the cross-links of **2B**, see Table 2). Second, the stoichiometry of **2A** differs from that of **2B** while the information is not available for **2C** and **2D**. One α-subunit and one β-subunit form a complex for **2A** [38], whereas **2B** is a tetramer with a homodimer of two β-subunits at the center [39]. Indeed, one pair of the cross-linked sites for **2B** (P0AGE9 residue 43 with P0A836 residue 172) requires the tetramer structure, whereas both pairs of cross-linked sites of **2A** are consistent with a dimeric complex structure. Consequently, we used the monomeric β-subunit as the docking input for **2A** and the dimeric β-subunit for **2B**. Based on the high sequence identities among **2B**, **2C,** and **2D** as well as the need of the dimeric β-subunit to rationalize some of the cross-links, we used the dimeric β-subunit structure as the input for docking **2C** and **2D**.

Group 3 consists of human (**3A**) and *E. coli* (**3B**) F1-ATP synthases. The bound structures show three α-subunits and three β-subunits that are close to the C_3_ symmetry but broken by the γ-subunit that binds to the center of the complex. The α and β-subunits have the same fold, but within each complex, the sequence identity is only 26% for both **3A** and **3B**. Between the complexes, the α-subunits have a sequence identity of 57% and the β-subunits 72%. Thus, we considered them distinct cases. We assumed that the synthases were stable without the γ-subunit and ignored the γ-subunit in docking and subsequent analysis. This is supported by the thermophilic *Bacillus* 1-ATP synthase, which has been crystallized both with and without a symmetry breaking subunit [40,41].

The remaining groups (4-8) involve complexes of two components with a single interface. Group 4 has a single member, the complex of human profilin-1 with β-actin (**4**), and is the only case for which all bound and unbound structures were available with high sequence identity (over 94%). This case is also an entry of the protein-protein docking benchmark that we maintain [6,42-45].

Group 5 contains a single complex, the heterodimer of human Alu binding proteins SRP9 and SRP14 bound to Alu RNA (**5**). We ignored the RNA component during docking assuming that the two proteins could form a complex without the RNA.

The single member of group 6 contains two interacting components (BamC and BamD) of the five-protein barrel assembly machinery (BAM) complex (**6**), responsible for the proper assembly of β-barrel proteins into the outer membrane of *E. coli*. BamC contains a 73-residue-long unstructured region essential for binding BamD (Figure 1) [46]. The unbound structure of the full-length BamC was not available, but even if it had been available, it might not have been suitable for rigid-body docking due to the unstructured region; consequently, we use the bound structure in our docking.

Group 7 is the complex of the GTPase Rab14 with a Rab GDP dissociation inhibitor (**7**). For the bound structure, we used the complex of the inhibitor with the prenylated YPT1 GTPase, having sequence identities of about 50% with the target.

Finally, group 8 consists of the human ElonginB-ElonginC complex (**8**) [47]. The bound structure includes the VHL tumor suppressor, but it was not included in the docking since VHL only contacts ElonginC and not ElonginB.

### Docking without cross-linking data

ZDOCK was used to predict complex structures for the twelve cases in the test set, using the component proteins as described above. We used interface RMSD between the predicted and bound structures (iRMSD) to assess the predictions. We applied an iRMSD cutoff of 5.0 Å to denote a prediction as a near-native structure or a ‘hit’ [48-50], except for test **2C**, **3A**, and **3B**, which had the interface RMSDs between the superposed and bound structures (I-RMSDs) ranging from 3 to 5 Å (Table 1). Since the I-RMSD forms a lower bound for the iRMSD, these cases required a cutoff of 7.5 Å to include at least one hit in the set of predictions. In case of distinct interfaces between the components (**1**, **3A**, and **3B)**, we assessed the docking predictions for each interface separately and also combined (claiming a prediction correct if either one of the interfaces observed in the bound structure matched the prediction). For **3A** and **3B**, whose bound complexes showed similar interfaces between the components with the differences caused by the symmetry breaking central chain, we based the assessment on the bound interface that yielded the lowest iRMSD.

Tables 3 and 4 show the results of unconstrained docking as well as docking combined with the various approaches of applying cross-linking constraints. Without constraints, most docking runs had a hit within the top 100 predictions (11 out of 15 interfaces), often within the top 10 (6 interfaces), but a top ranked hit was only found once. These results are in line with the ZDOCK performance on the protein-protein docking benchmark [6]. Interestingly, three of the four interfaces that had no hits within the top 100 predictions corresponded to the cases with multiple distinct interfaces (**1**, **3A**, and **3B**). We can speculate that the formation of the complex occurs in stages, in which the B:C (**1**) and A:D (**3A**, **3B**) interfaces are only stable after the B:A and A:E interfaces have formed. Alternatively, the chains that we ignored during docking (indicated in green in Figure 1) may be required for the complete complex formation.

**Table 3:** Docking results.

| Test Case<sup>1</sup> | Number of pairs of cross-linked sites | iRMSD cutoff used (Å) | Rank without constraints | Rank with Euclidean constraints (post-processing) |  |  | Rank with Euclidean constraints (within FFT) |  |  | Rank with void-volume constraints (post-processing) |  |  |
| --- | --- | --- | --- | --- | --- | --- | --- | --- | --- | --- | --- | --- |
|  |  |  |  | 30 Å | 35 Å | 40 Å | 30 Å | 35 Å | 40 Å | 35 Å | 40 Å | 45 Å |
| **1** | 2+3 | 5.0 | 6 | 1 | 1 | 3 | 1 | 1 | 3 | 1<br>(4th/75)<sup>2</sup> | 2<br>(3rd/58)<sup>2</sup> | 2 |
| **1 (B:A)** | 2 | 5.0 | 6 | 1 | 1 | 3 | 1 | 1 | 3 | 1<br>(4th/75)<sup>2</sup> | 2<br>(3rd/58)<sup>2</sup> | 2 |
| **1 (B:C)** | 3 | 5.0 | 164 | 4 | 5 | 7 | 4 | 6 | 7 | 15<br>(3rd/1033)<sup>2</sup> | 2 | 3 |
| **2A** | 2 | 5.0 | 3 | 1 | 1 | 1 | 1 | 1 | 1 | 1 | 1 | 1 |
| **2B** | 4<sup>3</sup> | 5.0 | 8 | none | 1 | 1 | 2 | 1 | 1 | none | 1 | 1 |
| **2C** | 1 | 7.5 | 200 | none | none | 824<br>(4th/1874)<sup>2</sup> | 2808 | 2483 | 988 | none | none | 617<br>(4th/1874)<sup>2</sup> |
| **2D** | 1 | 5.0 | 58 | none | none | 712<br>(4th/1332)<sup>2</sup> | none | none | 816 | none | none | 563<br>(4th/1332)<sup>2</sup> |
| **3A** | 2+5 | 7.5 | 11 | none | 1<br>(2nd/29)<sup>2</sup> | 1 | 1 | 1 | 1 | 1<br>(2nd/29)<sup>2</sup> | 1<br>(2nd/29)<sup>2</sup> | 1 |
| **3A (A:D)** | 2 | 7.5 | 469 | none | 3 | 8 | 1 | 1 | 9 | 2 | 3 | 7 |
| **3A (A:E)** | 5 | 7.5 | 11 | none | 1<br>(2nd/29)<sup>2</sup> | 1 | 1 | 1 | 1 | 1<br>(2nd/29)<sup>2</sup> | 1<br>(2nd/29)<sup>2</sup> | 1 |
| **3B** | 1 | 7.5 | 23 | 3 | 8 | 13 | 3 | 9 | 11 | 4 | 7 | 12 |
| **3B (A:D)** | 0 | 7.5 | 2020 | n/a<sup>4</sup> | n/a<sup>4</sup> | n/a<sup>4</sup> | n/a<sup>4</sup> | n/a<sup>4</sup> | n/a<sup>4</sup> | n/a<sup>4</sup> | n/a<sup>4</sup> | n/a<sup>4</sup> |
| **3B (A:E)** | 1 | 7.5 | 23 | 3 | 8 | 13 | 3 | 9 | 11 | 4 | 7 | 12 |
| **4** | 2 | 5.0 | 61 | 5 | 8 | 12 | 5 | 8 | 14 | 4 | 4 | 7 |
| **5** | 2 | 5.0 | 2 | 2 | 2 | 2 | 2 | 2 | 2 | 1 | 1 | 1 |
| **6** | 1 | 5.0 | 9 | none | none | 8<br>(2nd/17)<sup>2</sup> | none | none | 8 | none | none | 43<br>(3rd/141)<sup>2</sup> |
| **7** | 1 | 5.0 | 13 | 3 | 3 | 6 | 3 | 3 | 6 | 2 | 2 | 2 |
| **8** | 1 | 5.0 | 1 | 1 | 1 | 1 | 1 | 1 | 1 | 1 | 1 | 1 |

<sup>1</sup> When more than one distinct interface was present, the predictions were evaluated for the different interfaces combined, and separately for each specific interface as denoted in parentheses. See text for details.
<sup>2</sup> The top-ranked hit(s) prior to post-processing did not satisfy the cross-link(s), thus the top-ranked hit after filtering is not equivalent to the original top-ranked hit. In parentheses we show the number of the hit in the unfiltered list and its rank in the unfiltered list.
<sup>3</sup> Although there are seven observed pairs of cross-linked sites for case **2B** (Table 2), only four could be applied due to the incompleteness of the template used for homology modeling the unbound structure.
<sup>4</sup> There were no cross-linking constraints applicable to this interface.

**Table 4:** Number of test cases with hits in the 1, 5, and 10 highest ranked predictions (using the data from Table 3 and only the interface-specific evaluations for cases 1, 3A, and 3B)

| Number of predictions made | ZDOCK without constraints | ZDOCK with Euclidean constraints (post-processing) |  |  | ZDOCK with Euclidean constraints (within FFT) |  |  | ZDOCK with void-volume constraints (post-processing) |  |  |
| --- | --- | --- | --- | --- | --- | --- | --- | --- | --- | --- |
|  |  | 30 Å | 35 Å | 40 Å | 30 Å | 35 Å | 40 Å | 35 Å | 40 Å | 45 Å |
| 1 | 1 | 3 | 5 | 4 | 5 | 6 | 4 | 5 | 5 | 5 |
| 5 | 3 | 8 | 9 | 6 | 11 | 8 | 6 | 9 | 10 | 8 |
| 10 | 6 | 8 | 11 | 10 | 11 | 11 | 10 | 9 | 11 | 10 |

### Separating cross-links by interface

In all of our calculations, we assumed that the experimental cross-link data did not include false positives. Consequently, we applied hard cutoffs to the cross-link distances calculated for the predictions and required a prediction to satisfy all cross-links. It was straightforward to apply these requirements for binary complexes **2A** and **4**-**8**. Furthermore, **3B** has two distinct interfaces but only one cross-link, so we simply focused on the interface associated with the cross-link. For cases **1** and **3A** however, we had two distinct interfaces and multiple cross-links, and the cross-links for one interface may not be satisfied by the other interface of the same complex. Thus, it was essential that we assigned each cross-link to the appropriate interface. A similar issue arose for case **2B**, whereby cross-links could occur between the α-subunit and either chain of the β-subunit homodimer. In these situations, we need to group the cross-links by interface (**1** and **3A**) or chain (**2B**) so that we simultaneously apply only the cross-links that belong to the same interface or chain. Such grouping information, however, is not directly provided by the experimental cross-link data.

To group the cross-links by interface or chain, we assumed that two cross-links that are both satisfied by a docking prediction are more likely to be associated with the same interface (or chain) than with different interfaces. Thus, we defined a binary matrix with rows corresponding to the top 200 ZDOCK predictions, columns corresponding to the cross-links, and the elements set to 1 for the predictions that satisfy the cross-link (using Euclidean distance with a 30 Å cutoff) and 0 otherwise. We then calculated the correlation coefficients *r* between all pairs of columns to obtain a correlation matrix for each test case with each dimension corresponding to the total number of cross-links. Figure 2 depicts the six test cases with multiple cross-links, with the r>0.1 elements shaded. For cases **1, 2B**, and **3A**, a block-diagonal pattern arose, with each block corresponding to the cross-links that belong to the same interface or chain. As expected, the cross-links formed a single group for binary complexes **2A**, **4**, and **5**. Therefore, we applied each group of constraints separately during the docking runs. In addition to improving docking performance, the occurrence of multiple groups of cross-links provides information on the stoichiometry and topology of the complex, which we explored further to determine the symmetry of the complexes (see below).

**Figure 2:**
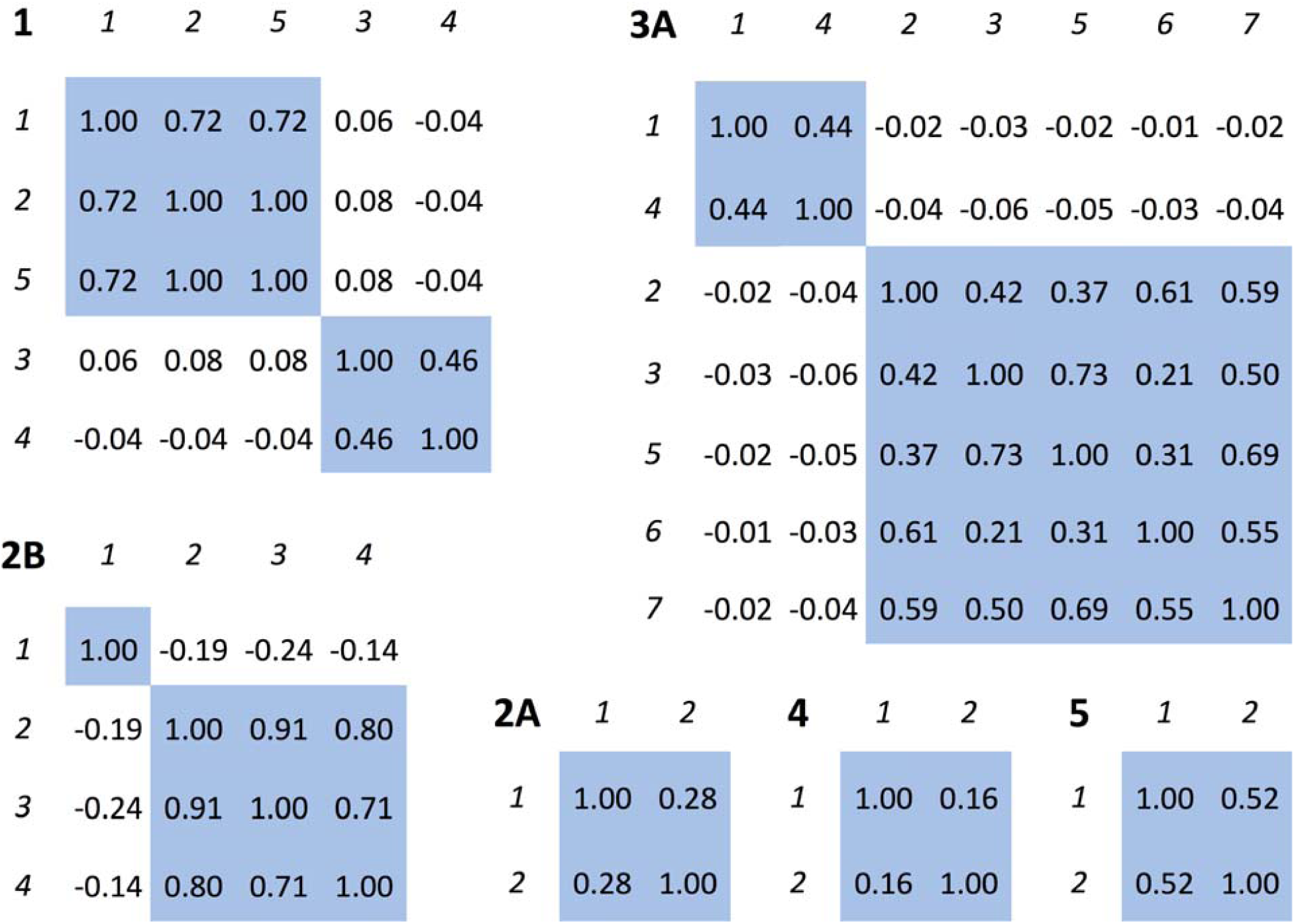
For the test cases with multiple cross-links, we calculated a binary vector for each cross-link with elements whether each of the top 200 ZDOCK predictions satisfied the cross-link (30 Å cutoff, Euclidean distance), and then computed the correlation coefficients (*r*) between all pairs of vectors. This figure shows the correlations *r* between the cross-links in each test case, with the positively correlated cross-links with values r>0.1 shaded. The indices correspond to the cross-links as ordered in Table 2, and were rearranged to obtain a block-diagonal shading pattern for cases **1** and **3A**. Each shaded block corresponds to a distinct interface.

### Filtering using Euclidean distances

We filtered out predictions with cross-linked sites further than 40 Å apart in Euclidean distance, which was based on the maximum observed distance of 39.0 Å in the bound structures (Table 2). Even with this loose cutoff, the highest ranking hit was eliminated for three docking runs—the second hit was retained for **6** and only the fourth hit was retained for **2C** and **2D**. As a result, incorporating cross-link data worsened the rank of the top hit for **2C** and **2D**. Nevertheless, the Euclidean filter resulted in four cases with a hit ranked as number one, a substantial improvement from just one case without the filter (Table 4).

When we tightened the cutoff to 35 Å, there were more cases for which the top-ranked hit did not pass the filter and several cases for which all hits were filtered out. However, the overall results improved compared with the 40 Å Euclidean filter or without filtering, judged by the number of cases with at least one hit in the top 1, top 5, or top 10 predictions (Table 4). Tightening the filter further to 30 Å, however, worsened the overall results. Thus, a 35 Å cutoff provided the Euclidean filter with the best balance for the present data set.

### Filtering using void-volume distances

Since void-volume distances for the bound structures were somewhat larger than Euclidean distances (Table 2), we increased the cutoff by 5 Å. Again, we observed a tradeoff between losing hits and improving the ranks of the retained hits, and we found the optimal cutoff to be 40 Å. The void-volume filter only performed slightly better than the Euclidean filter, increasing by one the total number of cases with a hit in the top 5 predictions, but leaving the other metrics unchanged. The results were similar to the 35 Å Euclidean filter at the level of individual test cases, with the ranks of the top hit no more than four positions apart.

### Euclidean constraints within FFT

The third type of constraint also uses Euclidean distances, but it restricts the translational search space of the docking algorithm and is therefore implemented within the fast Fourier transform (FFT) step of the ZDOCK algorithm. We tested the same cutoffs as for the Euclidean filter, and the performance was comparable with the above two post-processing approaches. The FFT-based constraint with a cutoff of 35 Å showed the best performance for the top 1 prediction, and with 30 Å for the top 5 predictions. For the top 10 predictions, it tied with the post-processing methods. Overall, the FFT-implemented constraint approach with a 30 Å cutoff yielded the best performance among all method and cutoff combinations. It is reasonable to consider the top 5 predictions in follow-up computational or experimental work, for which this approach had a success rate of 79% (11 out of 14 interfaces for which we had cross-links). None of the three tests cases for which this approach failed to generate a hit among the top 5 predictions (**2C**, **2D**, and **6**), was correctly predicted in the top 5 by the other method and cutoff combinations.

### Symmetry analysis

The occurrence of multiple groups of cross-links indicates that at least one component of the complex is a homo-multimer (as in the case of **2B**) or the components form multiple distinct interfaces. The latter can then lead to symmetric complexes (as in the case of **3A**) or a linear configuration (as in the case of **1**). To differentiate these three possibilities, we integrated the experimental cross-link data with docking to predict whether a complex had symmetry and whether this symmetry could be used to improve the docking performance.

Cases **1** and **3A** showed two distinct interfaces each (Figure 2), and we asked whether predictions for these interfaces could lead to symmetric complexes. We started with the top 5 predictions for each interface, obtained from the 30 Å FFT-implemented distance constraints. Combining these top 5 predictions yielded 25 predicted interface pairs. Starting from a monomer, we used each of these 25 interface pairs to sequentially add components, creating tetrameric (C_2_), hexameric (C_3_), and octameric (C_4_) complex structures for each interface pair. The resulting structures were not symmetric because the docking was performed without any symmetry information. We moved the component proteins as rigid bodies to reach a symmetric structure while keeping the deformation of the predicted interfaces to a minimum (Methods), and retained the symmetrized structure only if all the cross-link constraints were still satisfied and the iRMSD between the predicted interfaces and symmetrized interfaces did not exceed 2.5 Å. Figure 3 outlines the procedure for a single interface pair.

**Figure 3:**
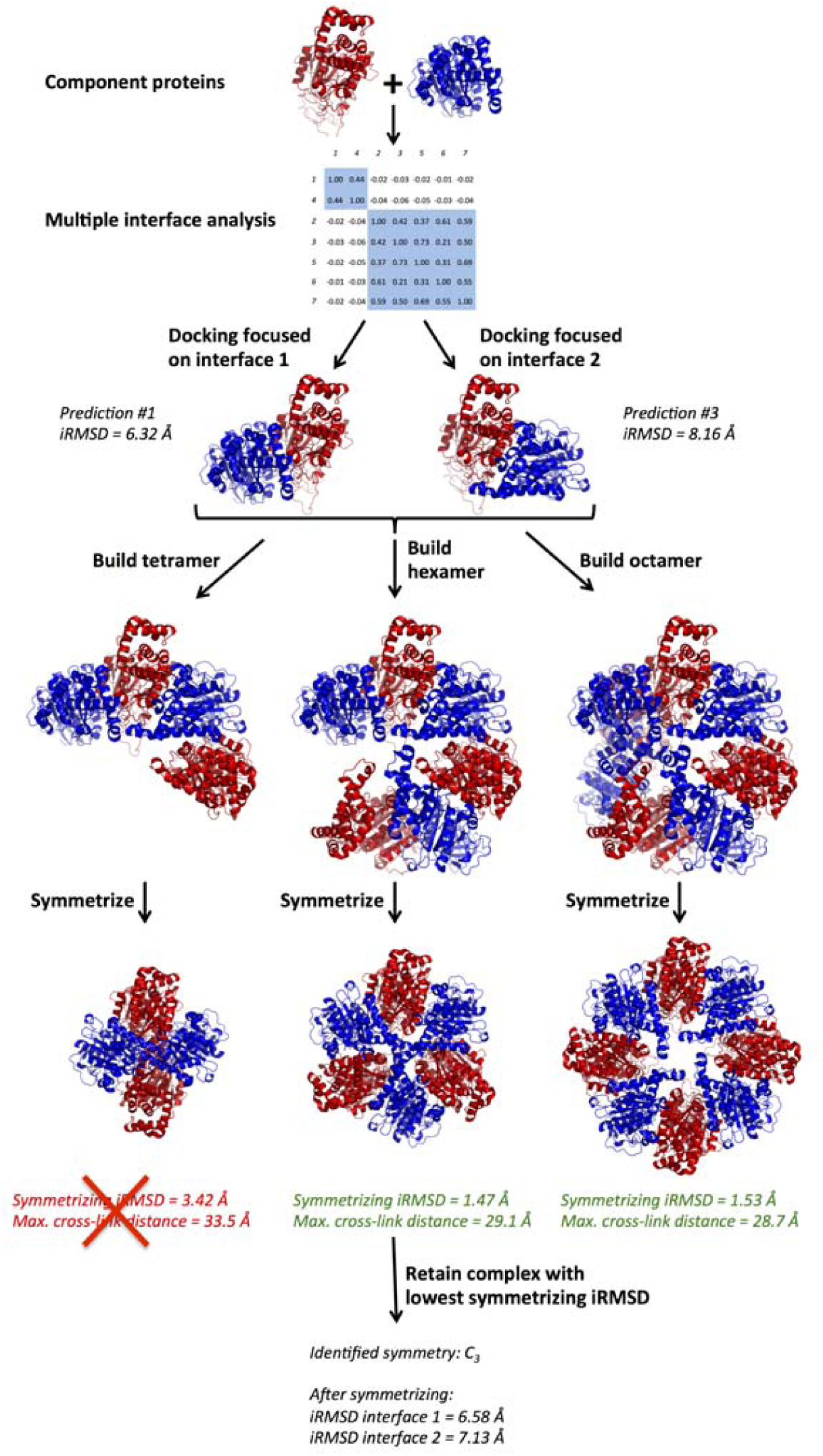
Example of the symmetry analysis (case **3A**, first and third prediction for interface A:E and A:D, respectively).

As shown in Tables S1 and S4, neither case **1** nor **3A** had a predicted interface pair that was consistent with the C_2_ symmetry. Case **1** only showed a single symmetry-consistent interface pair, which had the C_4_ symmetry (Tables S2 and S3). For **3A**, four predicted interface pairs were consistent with the C_3_ symmetry (Table S5), and five with C_4_ (Table S6). From the clash count (Tables S3 and S6) we see that a few octameric complexes symmetrize to unrealistic structures, but these examples also have starting interfaces that are far from the bound form. Table 5 integrates the results and they agree with the symmetries observed in the bound structures: we found seven symmetry-consistent pairs for **3A** (C_3_ in bound) and only one pair for **1** (non-symmetric in bound). Although the bound structure of **3A** is a hexamer, three of the seven interface pairs symmetrized to an octamer. Hexameric and octameric structures are difficult to distinguish by our algorithm because the difference between the C_3_ (hexamer) and C_4_ (octamer) symmetries is only 15 degrees per interface. In agreement with the bound structures, no predicted interface pairs yielded the C_2_ symmetry. Our results based on these two cases suggest that combining docking with cross-linking data can reveal whether a protein-protein complex is symmetric, although the predicted fold of the symmetry is not precise.

**Table 5:** iRMSDs before and after symmetrizing. The iRMSDs for symmetrized structures are shown only when they are close to the original predictions (symmetrizing iRMSD ≤ 2.5 Å and all cross-link distances ≤ 30 Å, see Tables S1-S6) and for the symmetry with the lowest symmetrizing iRMSD. Shaded iRMSD pairs are within same the cutoff as for a hit, defined as iRMSD ≤ 5.0 Å and 7.5 Å for case 1 and 3A, respectively.

| Interfaces |  | Case 1 |  |  |  |  | Case 3A |  |  |  |  |
| --- | --- | --- | --- | --- | --- | --- | --- | --- | --- | --- | --- |
| B:C/<br>A:D | B:A/<br>A:E | Original<br>iRMSD (Å) |  | Symmetrized<br>iRMSD (Å) |  |  | Original<br>iRMSD (Å) |  | Symmetrized<br>iRMSD (Å) |  |  |
|  |  | B:C | B:A | B:C | B:A | Sym | A:D | A:E | A:D | A:E | Sym |
| 1 | 1 | 24.07 | 3.59 |  |  |  | 6.32 | 5.72 | 6.60 | 5.93 | C<sub>4</sub> |
| 1 | 2 | 24.07 | 21.20 |  |  |  | 6.32 | 24.99 |  |  |  |
| 1 | 3 | 24.07 | 5.02 |  |  |  | 6.32 | 8.16 | 6.58 | 7.13 | C<sub>3</sub> |
| 1 | 4 | 24.07 | 3.51 |  |  |  | 6.32 | 17.76 |  |  |  |
| 1 | 5 | 24.07 | 3.76 |  |  |  | 6.32 | 14.48 |  |  |  |
| 2 | 1 | 18.27 | 3.59 |  |  |  | 9.80 | 5.72 | 9.26 | 6.43 | C<sub>4</sub> |
| 2 | 2 | 18.27 | 21.20 |  |  |  | 9.80 | 24.99 |  |  |  |
| 2 | 3 | 18.27 | 5.02 |  |  |  | 9.80 | 8.16 |  |  |  |
| 2 | 4 | 18.27 | 3.51 |  |  |  | 9.80 | 17.76 |  |  |  |
| 2 | 5 | 18.27 | 3.76 |  |  |  | 9.80 | 14.48 |  |  |  |
| 3 | 1 | 19.25 | 3.59 |  |  |  | 6.70 | 5.72 | 6.57 | 5.82 | C<sub>3</sub> |
| 3 | 2 | 19.25 | 21.20 |  |  |  | 6.70 | 24.99 |  |  |  |
| 3 | 3 | 19.25 | 5.02 |  |  |  | 6.70 | 8.16 | 7.09 | 7.65 | C<sub>3</sub> |
| 3 | 4 | 19.25 | 3.51 |  |  |  | 6.70 | 17.76 |  |  |  |
| 3 | 5 | 19.25 | 3.76 |  |  |  | 6.70 | 14.48 |  |  |  |
| 4 | 1 | 3.57 | 3.59 |  |  |  | 9.27 | 5.72 |  |  |  |
| 4 | 2 | 3.57 | 21.20 |  |  |  | 9.27 | 24.99 |  |  |  |
| 4 | 3 | 3.57 | 5.02 |  |  |  | 9.27 | 8.16 |  |  |  |
| 4 | 4 | 3.57 | 3.51 |  |  |  | 9.27 | 17.76 |  |  |  |
| 4 | 5 | 3.57 | 3.76 |  |  |  | 9.27 | 14.48 |  |  |  |
| 5 | 1 | 18.95 | 3.59 |  |  |  | 5.23 | 5.72 | 5.47 | 5.89 | C<sub>4</sub> |
| 5 | 2 | 18.95 | 21.20 | 20.24 | 20.70 | C<sub>4</sub> | 5.23 | 24.99 |  |  |  |
| 5 | 3 | 18.95 | 5.02 |  |  |  | 5.23 | 8.16 | 5.56 | 6.93 | C<sub>3</sub> |
| 5 | 4 | 18.95 | 3.51 |  |  |  | 5.23 | 17.76 |  |  |  |
| 5 | 5 | 18.95 | 3.76 |  |  |  | 5.23 | 14.48 |  |  |  |

Finally, we tested whether such a symmetry analysis could be used to improve the accuracy of complex structure prediction. For case **1** and **3A** combined, we considered 50 pairs of interface predictions, of which six showed both interfaces below the iRMSD cutoff for denoting a correct prediction (5.0 Å and 7.5 Å for cases **1** and **3A**, respectively), representing a success rate of 12%. Eight of the predicted interface pairs were symmetry-consistent, and five of the corresponding symmetrized structures had interfaces below the iRMSD cutoff (Table 5). Although the absolute number of correctly predicted interface pairs dropped from six to five, the success rate increased from 12% to 62.5% if we only retained the symmetry-consistent interface predictions.

## CONCLUSIONS

We demonstrated that incorporating cross-link data in *ab initio* protein-protein docking algorithms typically improve the rank of the first near-native prediction with a factor between two and ten. The success rates for the top 5 and top 10 predictions roughly doubled. Such improvements considerably increase the usefulness of protein-protein structure prediction algorithms. We tested several approaches to incorporate the cross-linking data in the docking calculations and found that using constraints in the translational search led to the best performance. Also, we showed that structures of symmetric complexes could be refined further, improving the predictions for the associated interfaces.

Our findings indicate that a distance cutoff of 30 Å as implemented in the FFT component of the docking algorithm yields the best overall performance, which is considerably shorter than the largest cross-link distances observed in the bound structures, close to 40 Å. Although the observed distances may be high due to differences in the complex structures between crystal and solution forms, it is possible that the performance for the few cases with large cross-link distances was sacrificed to improve the performance of the remaining test cases which represent the majority, leading to the best overall performance. For example, case **6** showed a large cross-link distance in the bound structure, and resulted in a hit with a similarly large constraint cutoff in the docking. Case **6**, however, was predicted incorrectly using the cutoff distances that showed the best performances overall.

Finally, our work suggests several directions for further algorithmic improvement. For example, we found that some correct predictions did not pass the cross-linking distance filters, even when the cutoffs were larger than the distances observed in the bound form. This may be due to the interface acting as a hinge, with small changes at the interface having large effects on the distal cross-linked sites. Thus, adjusting the cutoff distance according to the flexibility of the predicted interface may improve predictions. Similarly, we could assess the flexibilities of the regions of the proteins that are cross-linked, and adjust the cutoff distance accordingly. Alternatively, common structural refinement algorithms, which are often used as a post-processing step following the rigid-body docking, could be adapted to include cross-linking constraints

## MATERIALS AND METHODS

### Data sets

We compiled the cross-linking data sets from previous work and new experiments [29-34]. In all cases, cross-linking was performed using the BDP-NHP cross-linker described extensively in previous work [30]. Briefly, cross-linker was added to live cells resuspended in phosphate buffer (170 mM KH_2_PO_4_, pH 8.0), the cells were lysed, and protein lysates were digested with trypsin. Cross-linked peptide pairs were fractionated by strong cation exchange, enriched with monomeric avidin beads (Thermo) and loaded onto an in-house pulled 45cm C8 reverse phase column for injection into an LTQ-Velos FT-ICR instrument. ReACT [30] was run as previously described to identify cross-linked peptides in real time, and peptides were searched using SEQUEST.

### Protein-protein docking

We used our ZDOCK3.0 algorithm for the prediction of protein-protein complex structures [3,4,18-21]. ZDOCK inputs the structures of two constituent proteins and performs an exhaustive grid-based rigid-body search to predict their binary complex. The search returns an ensemble of predictions ranked using a scoring function, which includes shape complementarity, electrostatics, and the IFACE statistical potential [51,52]. Optimal results are typically obtained using X-ray crystallography structures as input, but also NMR, homology modeled, or *ab initio* modeled input structures can be utilized [53].

ZDOCK separates the full six-dimensional rigid-body space into a three-dimensional translational space and a three-dimensional rotational space. For each point in the rotational space, the best scoring translational pose is used as a prediction. In this study, we used 15° rotational sampling. Each docking run resulted in 3,600 predictions, which were ranked according to the docking scores.

### Cross-linking constraints

We considered three methods for incorporating cross-linking constraints in ZDOCK. The first two approaches involved post-processing, or filtering the set of predictions after the ZDOCK calculation. Cross-link distances were computed based on the predicted structures, and only when the distance was below a cutoff the constraint was considered satisfied, and the prediction retained. The two filtering approaches differed in the calculation of the cross-link distances. In the first, we used Euclidean (straight-line) distances between the C_β_ atoms. In the second, we used void-volume distances, computed with the command line version of the Xwalk program by Kahraman, MalmstrÖm and Aebersold (v0.6, using the C_β_ atom as anchor and the -bb flag in addition to the default options) [36]. Xwalk is grid-based and uses a breadth-first algorithm to obtain the shortest residue-residue distance that passes only through solvent accessible space. The third approach, FFT-based constraints, integrates the cross-linking constraints within the translational search of ZDOCK, following the algorithm presented by Xia [28]. In ZDOCK, Fast Fourier Transform (FFT) is used to generate a three-dimensional matrix that, for a point in the rotational space, contains the scores for all the (grid-based) translational coordinates. In a standard ZDOCK calculation, this matrix is then searched for the best score, and the corresponding complex structure is retained as a prediction. This structure may or may not satisfy the cross-links, hence the need for the filtering steps outlined above. In the FFT-constraint approach, on the other hand, we exploited the one-to-one correspondence between the score matrix elements and complex structures. For each score matrix element we could trivially compute its corresponding Euclidean cross-link distance, and when larger than the cutoff, marked the element as excluded. We repeated this procedure for all the cross-links observed for the complex. The subset of non-excluded elements was then scanned as usual, which yielded the best scoring complex structure that satisfied all the cross-links.

### Test set construction

We used BLAST [54], with a threshold of 30% sequence identity, to identify heteromeric complex structures in the PDB [35] for which we had cross-linking data available. We applied the following restrictions: First, the cross-linked sites needed to be part of the aligned regions. Second, the cross-linked sites needed to be resolved in the X-ray crystallography structure. The complexes were then investigated for suitability for protein-protein docking, using similar requirements as used for our protein-protein docking benchmarks [6]. For example, co-folded chains were excluded, as well as three-body (or higher order) interactions and protein-peptide like complexes. For the resulting complex list, we then searched the PDB for unbound structures. When finding bound or unbound structures that were less than 100% sequence identity with the cross-linked proteins, we used Modeller v.9.12 [54] to generate homology models, except for case **5** that had minimal sequence identity with the template, and yielded a more reasonable structure using I-TASSER [55-57]. We required at least one of the components to be available in its unbound form or as homology model based on an unbound template.

### Prediction assessment

To measure the quality of a prediction we used the C_α_ interface root-mean-square-deviation (iRMSD), which results from superposing the predicted interface onto the bound interface [6]. The interface includes all residues that have at least one atom within 10 Å of the binding partner in the bound structure. When the bound structure had multiple interfaces, we calculated the iRMSD for each interface separately and used the lowest value.

### Symmetry analysis

Starting with a single component and predictions for the two interfaces, we built multimeric structures by repeatedly adding components according to the predicted interfaces. We used PyMOL [58] for the superposition, and components were allowed to overlap if this followed from the interfaces. The resulting tetrameric, hexameric, and octameric structures were not symmetric because the predicted interfaces resulted from docking runs without any symmetry considerations. Therefore, we optimized each starting structure to the symmetric structure with the smallest deviation from the predicted interfaces, while keeping the components rigid. To achieve this, we used an optimization function that consisted of two components. The first component was the root mean square of the iRMSDs (defined above) between the predicted interfaces and the interfaces at the current optimization step. For the second component we followed the approach by Nilges [59]. We defined six centers for each of the two unique components (maximum and minimum along the three Cartesian axes) and calculated the distances between the paired centers located on different components. In a symmetric structure, the distances between centers across similar interfaces are identical. The optimization function thus contained the root mean square of the distance differences, which goes to zero at convergence. Although in principle the iRMSD component of the composite function is sufficient, we found that adding the specific symmetrizing component according to Nilges significantly improved the convergence behavior. For the optimization, we used steepest descent, with numerically calculated gradients. To speed up the optimizations we performed several thousand steps with only the interface similarity component, followed by additional steps using the full composite optimization function until a symmetric structure was obtained. The resulting function value, which is the root mean square of the iRMSDs (as the symmetrizing component was zero for the optimized structure) was denoted the *symmetrizing iRMSD* and used to measure how much the predicted interfaces deviated from symmetry.

## ACKNOWLEDGMENTS

This work was supported by NIH grants R01GM116960, R01GM086688, and U19AI107775.

## REFERENCES

[1] Wodak SJ, Vlasblom J, Turinsky AL, Pu S. Protein-protein interaction networks: the puzzling riches. Curr Opin Struc Biol 2013;23:941–53.

[2] Alber F, Förster F, Korkin D, Topf M, Sali A. Integrating diverse data for structure determination of macromolecular assemblies. Annu Rev Biochem 2008;77:443–77.

[3] Vreven T, Pierce BG, Hwang H, Weng Z. Performance of ZDOCK in CAPRI rounds 20-26. Proteins 2013;81:2175–82.

[4] Hwang H, Vreven T, Pierce BG, Hung J-H, Weng Z. Performance of ZDOCK and ZRANK in CAPRI rounds 13-19. Proteins 2010;78:3104–10.

[5] Lensink MF, Wodak SJ. Docking, scoring, and affinity prediction in CAPRI. Proteins 2013;81:2082–95.

[6] Vreven T, Moal IH, Vangone A, Pierce BG, Kastritis PL, Torchala M, et al. Updates to the Integrated Protein-Protein Interaction Benchmarks: Docking Benchmark Version 5 and Affinity Benchmark Version 2. J Mol Biol 2015;427:3031–41.

[7] Li L, Huang Y, Xiao Y. How to use not-always-reliable binding site information in protein-protein docking prediction. PLoS ONE 2013;8:e75936.

[8] van Ingen H, Bonvin AMJJ. Information-driven modeling of large macromolecular assemblies using NMR data. J Magn Reson 2014;241:103–14.

[9] Karaca E, Bonvin AMJJ. On the usefulness of ion-mobility mass spectrometry and SAXS data in scoring docking decoys. Acta Crystallogr D Biol Crystallogr 2013;69:683–94.

[10] Esquivel-Rodríguez J, Kihara D. Fitting multimeric protein complexes into electron microscopy maps using 3D Zernike descriptors. J Phys Chem B 2012;116:6854–61.

[11] Schneidman-Duhovny D, Rossi A, Avila-Sakar A, Kim SJ, Velázquez-Muriel J, Strop P, et al. A method for integrative structure determination of protein-protein complexes. Bioinformatics 2012;28:3282–9.

[12] Schmitz C, Bonvin AMJJ. Protein-protein HADDocking using exclusively pseudocontact shifts. J Biomol NMR 2011;50:263–6.

[13] Pons C, D’Abramo M, Svergun DI, Orozco M, Bernadó P, Fernandez-Recio J. Structural characterization of protein-protein complexes by integrating computational docking with small-angle scattering data. J Mol Biol 2010;403:217–30.

[14] Lasker K, Sali A, Wolfson HJ. Determining macromolecular assembly structures by molecular docking and fitting into an electron density map. Proteins 2010;78:3205–11.

[15] Ritchie DW, Kozakov D, Vajda S. Accelerating and focusing protein-protein docking correlations using multi-dimensional rotational FFT generating functions. Bioinformatics 2008;24:1865–73.

[16] Dominguez C, Boelens R, Bonvin A. HADDOCK: A protein-protein docking approach based on biochemical or biophysical information. J Am Chem Soc 2003;125:1731–7.

[17] Clore GM, Schwieters CD. Docking of protein-protein complexes on the basis of highly ambiguous intermolecular distance restraints derived from 1H/15N chemical shift mapping and backbone 15N-1H residual dipolar couplings using conjoined rigid body/torsion angle dynamics. J Am Chem Soc 2003;125:2902–12.

[18] Chen R, Li L, Weng Z. ZDOCK: an initial-stage protein-docking algorithm. Proteins 2003;52:80–7.

[19] Pierce BG, Wiehe K, Hwang H, Kim BH, Vreven T, Weng Z. ZDOCK server: interactive docking prediction of protein-protein complexes and symmetric multimers. Bioinformatics 2014;30:1771–3.

[20] Pierce BG, Hourai Y, Weng Z. Accelerating protein docking in lZDOCK using an advanced 3D convolution library. PLoS ONE 2011;6:e24657.

[21] Chen R, Weng Z. A novel shape complementarity scoring function for protein-protein docking. Proteins 2003;51:397–408.

[22] Herzog F, Kahraman A, Boehringer D, Mak R, Bracher A, Walzthoeni T, et al. Structural probing of a protein phosphatase 2A network by chemical cross-linking and mass spectrometry. Science 2012;337:1348–52.

[23] Rampler E, Stranzl T, Orban-Nemeth Z, Hollenstein DM, Hudecz O, Schloegelhofer P, et al. Comprehensive Cross-Linking Mass Spectrometry Reveals Parallel Orientation and Flexible Conformations of Plant HOP2-MND1. J Proteome Res 2015;14:5048–62.

[24] Doberenz C, Zorn M, Falke D, Nannemann D, Hunger D, Beyer L, et al. Pyruvate formate-lyase interacts directly with the formate channel FocA to regulate formate translocation. J Mol Biol 2014;426:2827–39.

[25] Schweppe DK, Chavez JD, Lee CF, Caudal A, Kruse SE, Stuppard R, et al. Mitochondrial protein interactome elucidated by chemical cross-linking mass spectrometry. Proceedings of the National Academy of Sciences 2017;114:1732–7.

[26] van Zundert GCP, Bonvin AMJJ. DisVis: quantifying and visualizing accessible interaction space of distance-restrained biomolecular complexes. Bioinformatics 2015;31:3222–4.

[27] Kahraman A, Herzog F, Leitner A, Rosenberger G, Aebersold R, Malmström L. Cross-link guided molecular modeling with ROSETTA. PLoS ONE 2013;8:e73411.

[28] Xia B, Vajda S, Kozakov D. Accounting for pairwise distance restraints in FFT-based protein-protein docking. Bioinformatics 2016.

[29] Zheng C, Weisbrod CR, Chavez JD, Eng JK, Sharma V, Wu X, et al. XLink-DB: database and software tools for storing and visualizing protein interaction topology data. J Proteome Res 2013;12:1989–95.

[30] Weisbrod CR, Chavez JD, Eng JK, Yang L, Zheng C, Bruce JE. In vivo protein interaction network identified with a novel real-time cross-linked peptide identification strategy. J Proteome Res 2013;12:1569–79.

[31] Chavez JD, Schweppe DK, Eng JK, Zheng C, Taipale A, Zhang Y, et al. Quantitative interactome analysis reveals a chemoresistant edgotype. Nat Commun 2015;6:7928.

[32] Schweppe DK, Harding C, Chavez JD, Wu X, Ramage E, Singh PK, et al. Host-Microbe Protein Interactions during Bacterial Infection. Chem Biol 2015;22:1521–30.

[33] Navare AT, Chavez JD, Zheng C, Weisbrod CR, Eng JK, Siehnel R, et al. Probing the protein interaction network of Pseudomonas aeruginosa cells by chemical cross-linking mass spectrometry. Structure 2015;23:762–73.

[34] Chavez JD, Schweppe DK, Eng JK, Bruce JE. In Vivo Conformational Dynamics of Hsp90 and Its Interactors. Cell Chem Biol 2016;23:716–26.

[35] Berman HM. The Protein Data Bank. Nucleic Acids Research 2000;28:235–42.

[36] Kahraman A, Malmström L, Aebersold R. Xwalk: computing and visualizing distances in cross-linking experiments. Bioinformatics 2011;27:2163–4.

[37] Maderna A, Doroski M, Subramanyam C, Porte A, Leverett CA, Vetelino BC, et al. Discovery of cytotoxic dolastatin 10 analogues with N-terminal modifications. J Med Chem 2014;57:10527–43.

[38] Fraser ME, James MN, Bridger WA, Wolodko WT. Phosphorylated and dephosphorylated structures of pig heart, GTP-specific succinyl-CoA synthetase. J Mol Biol 2000;299:1325–39.

[39] Wolodko WT, Fraser ME, James MN, Bridger WA. The crystal structure of succinyl-CoA synthetase from Escherichia coli at 2.5-A resolution. J Biol Chem 1994;269:10883–90.

[40] Shirakihara Y, Leslie AG, Abrahams JP, Walker JE, Ueda T, Sekimoto Y, et al. The crystal structure of the nucleotide-free alpha 3 beta 3 subcomplex of F1-ATPase from the thermophilic Bacillus PS3 is a symmetric trimer. Structure 1997;5:825–36.

[41] Shirakihara Y, Shiratori A, Tanikawa H, Nakasako M, Yoshida M, Suzuki T. Structure of a thermophilic F1-ATPase inhibited by an ε-subunit: deeper insight into the ε-inhibition mechanism. FEBS J 2015;282:2895–913.

[42] Chen R, Mintseris J, Janin J, Weng Z. A protein-protein docking benchmark. Proteins 2003;52:88–91.

[43] Mintseris J, Wiehe K, Pierce B, Anderson R, Chen R, Janin J, et al. Protein-Protein Docking Benchmark 2.0: an update. Proteins 2005;60:214–6.

[44] Hwang H, Pierce B, Mintseris J, Janin J, Weng Z. Protein-protein docking benchmark version 3.0. Proteins 2008;73:705–9.

[45] Hwang H, Vreven T, Janin J, Weng Z. Protein-protein docking benchmark version 4.0. Proteins 2010;78:3111–4.

[46] Kim KH, Aulakh S, Paetzel M. Crystal structure of β-barrel assembly machinery BamCD protein complex. J Biol Chem 2011;286:39116–21.

[47] Stebbins CE, Kaelin WG, Pavletich NP. Structure of the VHL-ElonginC-ElonginB complex: implications for VHL tumor suppressor function. Science 1999;284:455–61.

[48] Vreven T, Hwang H, Weng Z. Integrating atom-based and residue-based scoring functions for protein-protein docking. Protein Sci 2011;20:1576–86.

[49] Pierce B, Weng Z. ZRANK: reranking protein docking predictions with an optimized energy function. Proteins 2007;67:1078–86.

[50] Vreven T, Hwang H, Weng Z. Exploring angular distance in protein-protein docking algorithms. PLoS ONE 2013;8:e56645.

[51] Mintseris J, Pierce B, Wiehe K, Anderson R, Chen R, Weng Z. Integrating statistical pair potentials into protein complex prediction. Proteins 2007;69:511–20.

[52] Mintseris J, Weng Z. Optimizing protein representations with information theory. Genome Inform 2004;15:160–9.

[53] Rodrigues JPGLM, Melquiond ASJ, Karaca E, Trellet M, van Dijk M, van Zundert GCP, et al. Defining the limits of homology modeling in information-driven protein docking. Proteins 2013;81:2119–28.

[54] Sali A, Blundell TL. Comparative protein modelling by satisfaction of spatial restraints. J Mol Biol 1993;234:779–815.

[55] Roy A, Kucukural A, Zhang Y. I-TASSER: a unified platform for automated protein structure and function prediction. Nat Protoc 2010;5:725–38.

[56] Zhang Y. I-TASSER server for protein 3D structure prediction. BMC Bioinformatics 2008;9:40.

[57] Yang J, Yan R, Roy A, Xu D, Poisson J, Zhang Y. The I-TASSER Suite: protein structure and function prediction. Nat Methods 2015;12:7–8.

[58] The PyMOL Molecular Graphics System, Version 1.8 Schrödinger, LLC. n.d.

[59] Nilges M. A calculation strategy for the structure determination of symmetric dimers by 1H NMR. Proteins 1993;17:297–309.

